# Proteome-aware organ proxy aging clocks

**DOI:** 10.64898/2026.04.24.720503

**Authors:** Hao Xu, Jiawei Chen, Dongxu Chen, Kehang Mao, Jing-Dong J. Han

**Affiliations:** Peking-Tsinghua Center for Life Sciences, Academy for Advanced Interdisciplinary Studies, Center for Quantitative Biology (CQB), Peking University, Beijing 100871, China; Peking University Chengdu Academy for Advanced Interdisciplinary Biotechnologies, Chengdu, Sichuan 610213, China

## Abstract

The development of minimally invasive multi-organ aging clocks, established through the deconvolution of plasma proteomics, has provided a convenient tool to assess the organ heterogeneity of aging. However, prior studies relied on bulk transcriptomic data for organ marker identification, which may lead to the potential misidentification of protein markers, and their research scope was largely confined to a few common diseases. To address these limitations, this study integrated multi-dimensional data to refine organ-enriched marker panels by incorporating organ-specific proteome information, and developed Proteome-Aware Organ Proxy Proteome Aging Clock (PAOPAC). PAOPAC exhibited decelerated biological age corresponding to improved physiological phenotypes across two independent external datasets, demonstrating its generalizability. We then leveraged PAOPAC to generate a comprehensive disease-aging landscape and to investigate the process of chronological and biological aging. Our analyses revealed that the majority of diseases are associated with an accelerated aging phenotype.

## Introduction

Accurately quantifying the biological aging process is highly dependent on reliable aging clocks. However, identifying clinically valuable aging biomarkers and constructing universally applicable aging clocks has long been a core unsolved problem in this field^1,2^ and presents multiple methodological challenges. Firstly, regarding the choice of measuring omics, although phenomics (e.g., imaging, physiological indicators) offers advantages of being non-invasive or minimally invasive, the features they capture are often downstream outputs of aging, making it difficult to deeply parse their upstream molecular mechanisms without other sequencing-based omics^3,4^. While transcriptomics and epigenomics can provide rich mechanistic insights, the *in vivo*, longitudinal, and repeated sampling from specific organs (such as the brain, heart, liver) poses significant ethical and technical hurdles^5^. Secondly, there is an inherent trade-off between the generalizability and specificity of aging clocks. Most existing clocks are built using single biological sample and tailored to specific tissues, resulting in a lack of cross-tissue universality^6,7^. Conversely, clocks that integrate multi-organ shared features typically sacrifice organ-level specificity, hindering the resolution of organ-specific heterogeneity in aging^5^.

Fortunately, plasma proteomics demonstrates unique integrative potential. Regarding sample accessibility, plasma samples can be obtained minimally invasively via routine venipuncture, facilitating longitudinal collection and repeated measurements in large-scale population cohorts. Furthermore, besides the proteins with a functional role in plasma, there are also tissue actively secrete or leakage proteins in the circulation^8^. Theoretically, computational methods can deconvolve specific signals from different organ systems. And both work in experimental animals and human demonstrate that the abundance of tissue leakage proteins in plasma can reflect physiological or pathological states of their tissue of origin^9,10^. Finally, on the technological front, high-throughput proteomics platforms based on affinity reagents, such as Olink and SomaScan, are increasingly mature, enabling the systematic detection of hundreds to thousands of plasma proteins in large cohorts^11,12^. Despite the breakthroughs in current research utilizing the plasma proteome, they still face several key limitations that warrant in-depth investigation.

First, the vast majority of current aging studies rely on data generated from a single technology platform^13-16^. This dependency may introduce platform-specific technical biases, limiting the generalizability of the conclusion. Second, currently developed clocks that deconvolve organ-proxy aging signals from the plasma proteome are typically based on GTEx (Genotype-Tissue Expression) dataset^14-16^, which is a measurement at bulk-RNA level^17^. However, plasma protein abundance is regulated not only by transcription but also by a multitude of complex biological processes, including post-transcriptional, translational and post-translational regulations^18-20^. Third, current studies exploring associations between plasma proteomic aging clocks and diseases often focus on a few typical age-related diseases (e.g., Alzheimer’s disease, type 2 diabetes). This selective focus overlooks a systematic investigation of a broader disease spectrum, lacking research on non-canonical age-related diseases or comorbidity patterns, which limits our understanding of aging as a common risk factor within a complex network of multiple diseases.

To address the summarized limitations, this study utilized plasma proteomic data from over 50,000 individuals in the UK Biobank (UKB) based on the Olink platform^21^ as the primary discovery cohort, and another independent Olink dataset as external validation, alongside a combined cohort of over 300 individuals based on the SomaScan platform^16,22,23^ to cross-examine the conclusions found in discovery cohort. Comparative analyses on aging-related proteins, their temporal change patterns, and associated pathways were conducted to derive platform-agnostic insights into aging. This study also incorporated the Human Protein Atlas (HPA)^24^ and Cell Marker 2.0 (CM2)^25^ databases, providing complementary evidence at the bulk proteome and single-cell transcriptome levels, respectively. This approach enhances the theoretical foundation for constructing Multi-Organ Proteome Aging Clock (PAOPAC). A comprehensive comparison between ICD-10 coded event outcomes and organ aging rates was performed to construct a complete disease-aging landscape.

## Results

### More accurate organotypic proteins reveal age-dependent dynamics

We utilized proteomic data from the UKB^21^ as the discovery dataset (Supplementary Fig. 1a). Furthermore, we cross-examined three smaller public datasets based on the aptamer-based SomaScan platform^16,22,23^ (Methods, Fig. 1a and Supplementary Fig. 1a) for protein-level and pathway-level consistency.

**Figure 1.**
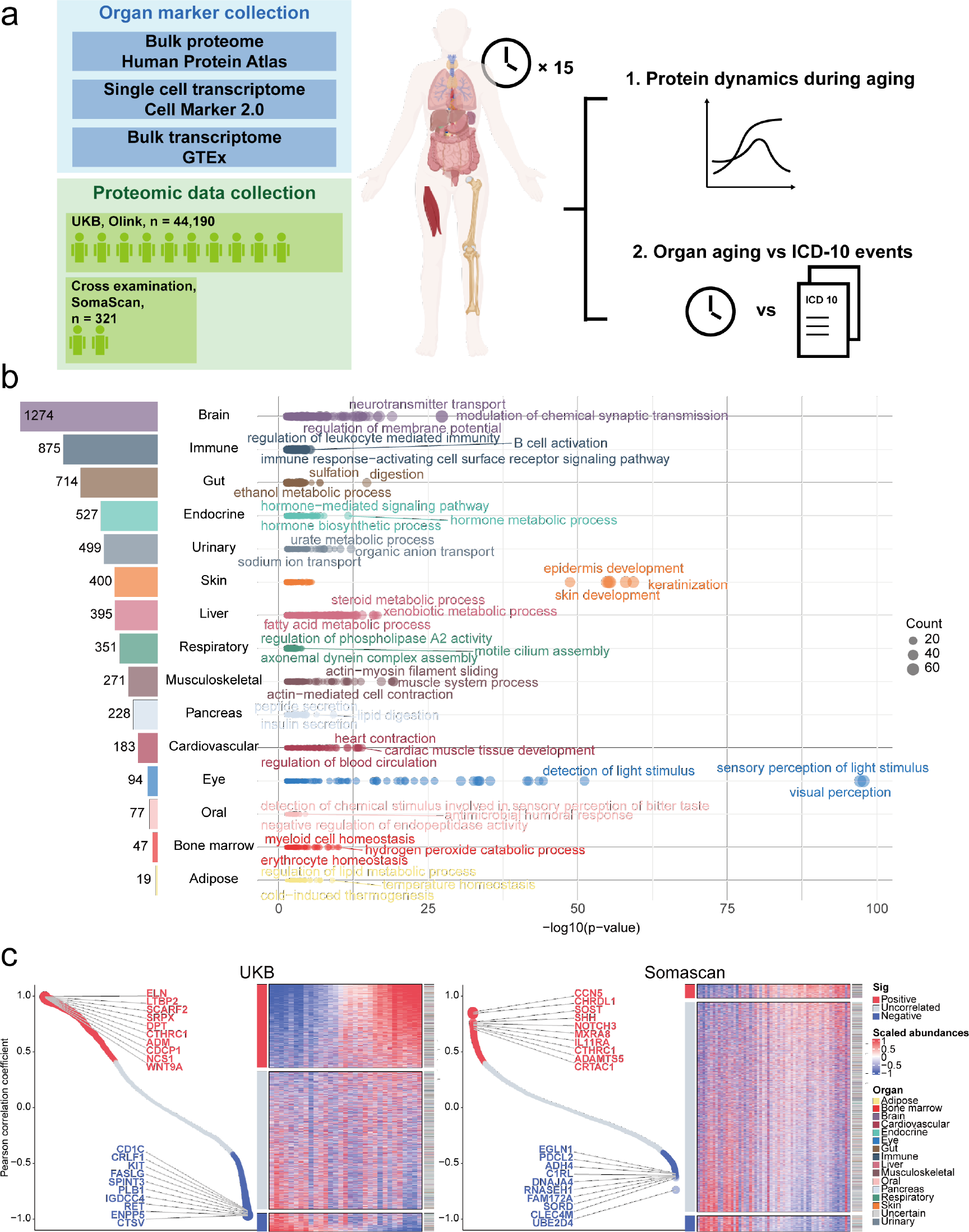
Study design, organ-enriched markers identification and their correlation with aging. **a**. Three dimensions of data (bulk proteome, single cell transcriptome and bulk transcriptome) were combined to identify organ-enriched markers, based on which organ-proxy aging clocks were constructed. 14 organ-proxy clocks and one conventional clock were developed to generate a comprehensive disease-aging landscape. **b**. A curated list of markers covering 15 organs/tissues was presented, with each organ-enriched marker set showing significant enrichment in physiological pathways corresponding to their respective organs. **c**. Proteins showing linear association with age in two platforms.

As noted earlier, organ enriched markers for current organ aging clocks were primarily derived from the GTEx dataset^14-16^, which reflects bulk RNA-level measurements^17^. However, due to post-transcriptional, translational and post-translational regulation, protein abundance does not always correlate with RNA expression^18-20^. To address this discrepancy, we prepared three marker sets representing different molecular levels: the Human Protein Atlas (HPA) for bulk proteome, CellMarker 2.0 (CM2) for single-cell transcriptome, and the GTEx dataset for bulk transcriptome^17,24,25^ (Methods and Fig. 1a). An initial analysis revealed that only a small proportion of organ markers used in existing clocks were consistent with HPA data, while at least 20% of proteins showed high expression in other organs (Supplementary Fig. 1b). But we found that markers derived solely from GTEx constitute approximately 20% of the final usable proteins in both platforms (Supplementary Fig. 2a). So, we decided to use this bulk-transcriptome data as a complementary resource after a stringent filter. Ultimately, we integrated a final marker list encompassing 15 tissues/organs and 5,954 proteins (Fig. 1b).

To further validate these organ enriched markers are associated with the organ functions, we performed GO term enrichment analysis. The results indicated strong associations between the marker sets and relevant organ functions. For example, brain markers were enriched for terms such as modulation of chemical synaptic transmission, neurotransmitter transport, and regulation of membrane potential. Even for organs with relatively few markers, such as bone marrow, functionally relevant terms like myeloid cell homeostasis and erythrocyte homeostasis were significantly enriched (Fig. 1b).

We next investigated proteins exhibiting age-dependent dynamics across platforms and traced their tissue origins. Linear age-associated proteins were identified by calculating Pearson correlation coefficients (PCC) between expression levels and age. We observed that ELN, SCARF2, DPT and SOST were positively correlated with age (Fig. 1c). Among these, ELN and SCARF2 are core components of a recently published proteomic aging clock^26^. Also, ELN and its fragment can reduce healthspan and lifespan^27^. DPT has been shown to ameliorate inflammation in adipose tissue^28^, and interacts with TGF-β, potentiating its effects. Notably, TGF-β is a key mediator of the senescence-associated secretory phenotype (SASP) and promotes systemic aging^29^. SOST is upregulated in calcifying aortic valve disease^30^. Proteins that decreased with age (Fig. 1c) like IGDCC4, KIT and UBE2D4, have been identified as a protective factor against dementia^31^, and age-related non-alcoholic steatohepatitis (NASH)^32^, or helps maintain proteome homeostasis in youth^33^.

We noted a substantial number of age-uncorrelated proteins in all datasets, particularly in SomaScan, likely due to its broader target coverage. To capture more nuanced temporal patterns, we applied the Fuzzy C-Means (FCM) clustering algorithm^34^ to group proteins with similar expression trajectories over time. Four distinct expression patterns were identified in both datasets: ‘Up-Down’, ‘Up’, ‘Down’, and ‘Down-Up’ (Fig. 2a and Fig. 2b). We found the overlap of specific proteins between platforms was limited, and SomaScan detected more down-regulated proteins, possibly owing to its higher sensitivity (Fig. 2c), a finding consistent with previous studies^35-37^. Given that expression patterns were conserved across platforms, we hypothesized that broader functional insights might reconcile technical differences. Indeed, pathway enrichment analysis revealed relatively more similarity at the functional level (Fig. 2c). These consistent pathways include the ‘Up-Down’ pattern enriched leukocyte-related pathways and negative regulation of proteolysis (Fig. 2a and Fig. 2b); the ‘Up’ pattern enriched extracellular matrix organization and inflammation-related pathways (Fig. 2a and Fig. 2b), both established hallmarks of aging^38^; the ‘Down’ pattern enriched PI3K-Akt signaling pathway, which enhances mitochondrial function^39^, (Fig. 2a and Fig. 2b) and may contribute to increased telomere vulnerability^40^; and finally the ‘Down-Up’ pattern enriched vesicle organization and hemostasis (Fig. 2a and Fig. 2b). Tissue-origin analysis further indicated an age-related reduction in brain-derived proteins in plasma (Fig. 2a and Fig. 2b).

**Figure 2.**
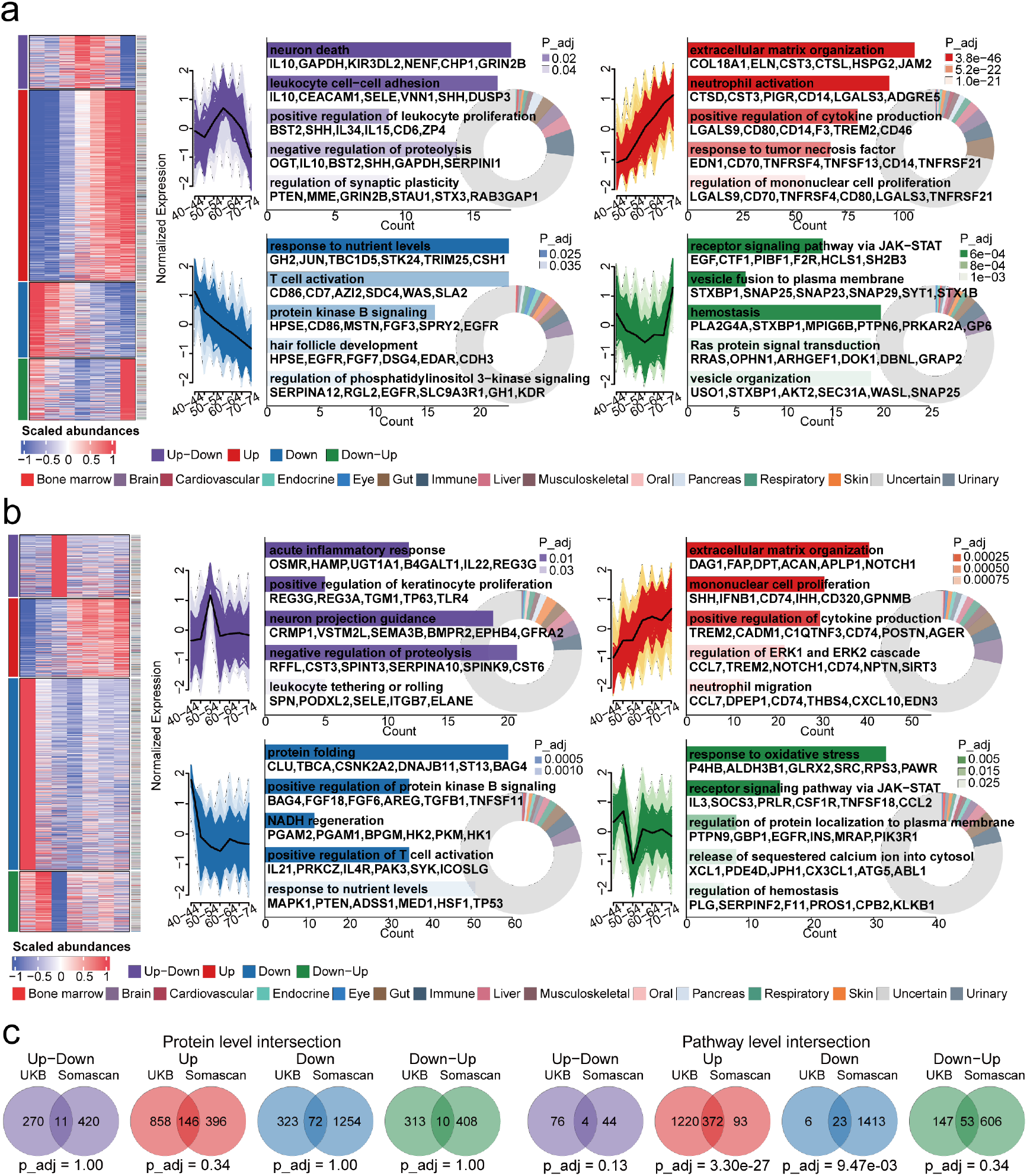
Age-related protein dynamics and cross-platform comparison. **a-b**. Four distinct proteomic dynamic change patterns were identified in the UKB cohort (a) and SomaScan cohort (b), along with the enriched pathways and the proportion of tissue-enriched proteins under each pattern. **c**. Aging-related proteins are different but their enriched pathways are relatively more conserved across platforms. P value calculated by hypergeometric test, and adjusted by BH.

In summary, while differences exist at the protein level between platforms, we observed relatively more consistent dynamic patterns and associated pathways at a broader, functional level. This macro-level analysis may help reconcile platform-specific discrepancies and offers insights for future research integrating data from different proteomic platforms.

### Construction of more accurate organ-proxy aging clocks

To quantify the heterogeneity of aging rates across different organs, we constructed organ-proxy aging clocks based on the plasma proteome, following the approach of previous studies^14-16^, but using the marker list identified in this work. We evaluated seven machine learning methods, among which LGBM and XGBoost demonstrated superior performance, particularly for organs with fewer markers (Fig. 3a). Due to its faster training speed (∼32 hours vs. ∼50.5 hours for XGBoost), LGBM was selected as the final model for building the aging clocks. During model training, covariates were included for adjustment, and the Boruta algorithm was applied for feature selection (Methods). We ultimately developed one conventional aging clock trained on all plasma proteins and 14 organ-proxy clocks. On the held-out validation set (20% of the total data), 12 of the organ clocks achieved a Pearson correlation coefficient (PCC) above 0.4, with the exceptions being the Eye, Bone Marrow, and Oral clocks (Fig. 3b).

**Figure 3.**
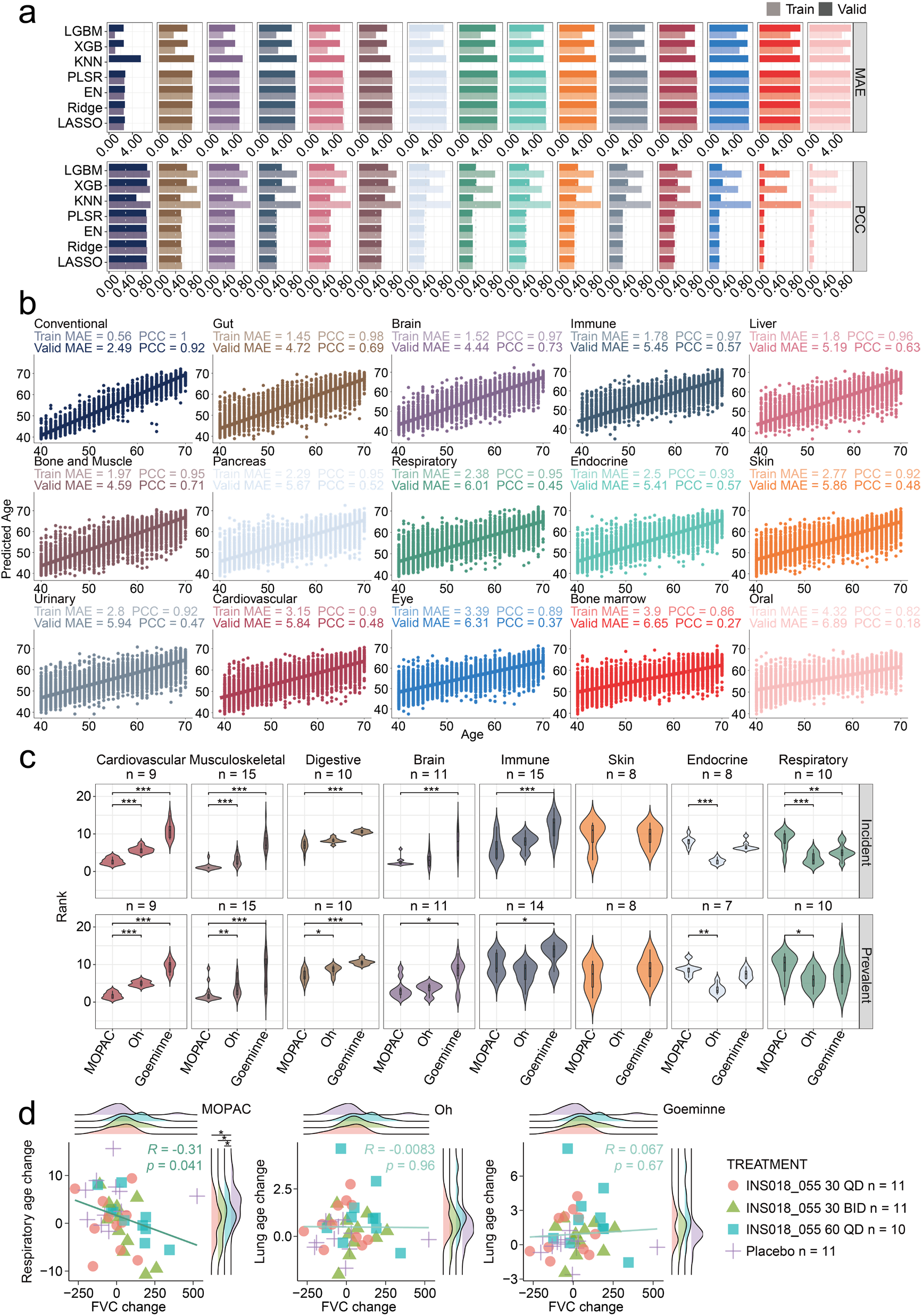
Construction and comparison of organ proxy aging clocks. **a**. Among the seven machine learning methods evaluated, LGBM demonstrated the best performance, particularly for organs with a limited number of available protein markers. **b**. Fourteen organ-proxy aging clocks and one conventional clock were constructed using the LGBM algorithm. **c**. Comparison of organ proxy performance between our clocks and two previously published work. PAOPAC performed particularly well for Cardiovascular, Musculoskeletal, Digestive and Brain, while remaining competitive for Immune, Skin, Endocrine and Respiratory (Wilcox test). **d**. Performance evaluation of our clocks alongside two published clocks on an external IPF dataset. Only the respiratory clock from this study reflected the post-treatment improvement in FVC was negatively associated with the reversal of physiological age (t-test). UKB, UK Biobank. FVC, forced vital capacity. For panels d and e, non-label: no significance, *: 0.01 < p < 0.05, **: 0.001 < p < 0.01, ***: p < 0.001.

Given that our Conventional clock was trained on the full proteome-consistent with previous studies-we further benchmarked its characteristics against prior models. Using Elastic Net (EN), Goeminne *et al*.^15^ retained the largest number of features, followed by Oh *et al*.^14^, who utilized LASSO regression. In contrast, our model employed LGBM coupled with the Boruta algorithm for rigorous feature selection, resulting in the most parsimonious feature set. Notably, nearly all of the markers identified by our approach constituted a subset of those identified by the other two clocks (Supplementary Fig. 3a). This refined marker selection might have contributed to the enhanced generalizability to our approach.

We next compared our organ-proxy aging clocks with previously published models. As the initial study by Oh *et al*. used the SomaScan platform^16^, making direct comparisons challenging, we focused on more recent work by Goeminne *et al*. and Oh *et al*.^14,15^. Although these studies also relied on the UKB proteomic data, the discrepancies in data splitting protocols make direct benchmark comparisons (e.g., MAE) problematic due to the unverifiable risk of data leakage.

However, a robust aging clock should also effectively capture variations in biological age, such as functional decline under disease conditions. To further examine the ability of these clocks to detect biological aging differences, we leveraged detailed ICD-10 diagnostic records available in the UKB to evaluate associations between organ aging states and related diseases.

First, we calculated the age-adjusted biological age difference for each organ-proxy clock (Organ-cAgeDiff, see Methods). Leveraging ICD-10 chapter codes, we mapped specific diseases to their corresponding target organs. The Conventional was excluded, because of its lacking of a specific anatomical target. Then, for both prevalent and incident cases, we employed Logistic or Cox proportional hazards regression models to determine the Odds Ratios (OR) or Hazard Ratios (HR) for each organ-proxy clocks in relation to these diseases. Within each disease category, the OR/HR values of all organs were ranked in descending order. A rank closer to 1 indicates that the corresponding organ-proxy clock possesses superior diagnostic or predictive power for diseases within that anatomical domain compared to other organ clocks, thereby validating its effectiveness as an organ-level biological proxy. PAOPAC performed particularly well for Cardiovascular, Musculoskeletal, Digestive and Brain, while remaining competitive for Immune, Skin, Endocrine and Respiratory (Fig. 3c). Although PAOPAC included two novel organs: Eye and Bone marrow, they didn’t show good organ-proxy performance, same for the Urinary clock in comparison (Supplementary Fig. 3b). Thus, these three organs will be excluded in future research.

Next, we further evaluated the ability of different clocks to discriminate intervention effects. We tested clocks using an external dataset comprising plasma samples from 43 patients with idiopathic pulmonary fibrosis (IPF) collected before and after 12 weeks of treatment^41^. Notably, only the Respiratory clock in PAOPAC successfully detected a significant reduction in biological age within the treatment group compared to the placebo (Supplementary Fig. 3c). This shift likely reflects a meaningful improvement in pulmonary physiological integrity. Supporting this interpretation, we observed that reductions in PAOPAC Respiratory age were significantly correlated with improvements in forced vital capacity (FVC), which were undetectable by the Goeminne or Oh Lung clock (Fig. 3d), thereby underscoring the clinical relevance and sensitivity of our organ-proxy model.

In summary, by leveraging organ-enriched proteomic markers, rather than relying solely on transcriptomic data, our clocks demonstrated improved organ-proxy performance and clinical application potential.

### A comprehensive disease-aging relation landscape

While previous studies have primarily focused on a limited set of common diseases-such as type 2 diabetes, heart failure, and Alzheimer’s disease--a comprehensive analysis of the relationship between disease and organ aging has been lacking. In this study, we therefore compared Organ-cAgeDiff between patients and healthy controls across 101 prevalent disease groups and 104 incident disease groups. Our analysis reveals that most disease categories are not limited to the aging of a single organ but are associated with accelerated aging across multiple organ-proxies (OR/HR > 1, indicated by red shading in Fig. 4a). And, Organ-cAgeDiff exhibited a high degree of consistency between prevalent cases and incident risks (Fig. 4b). This suggests that the Organ-cAgeDiff not only reflects current pathological states but also possesses prospective predictive value for future disease onset.

**Figure 4.**
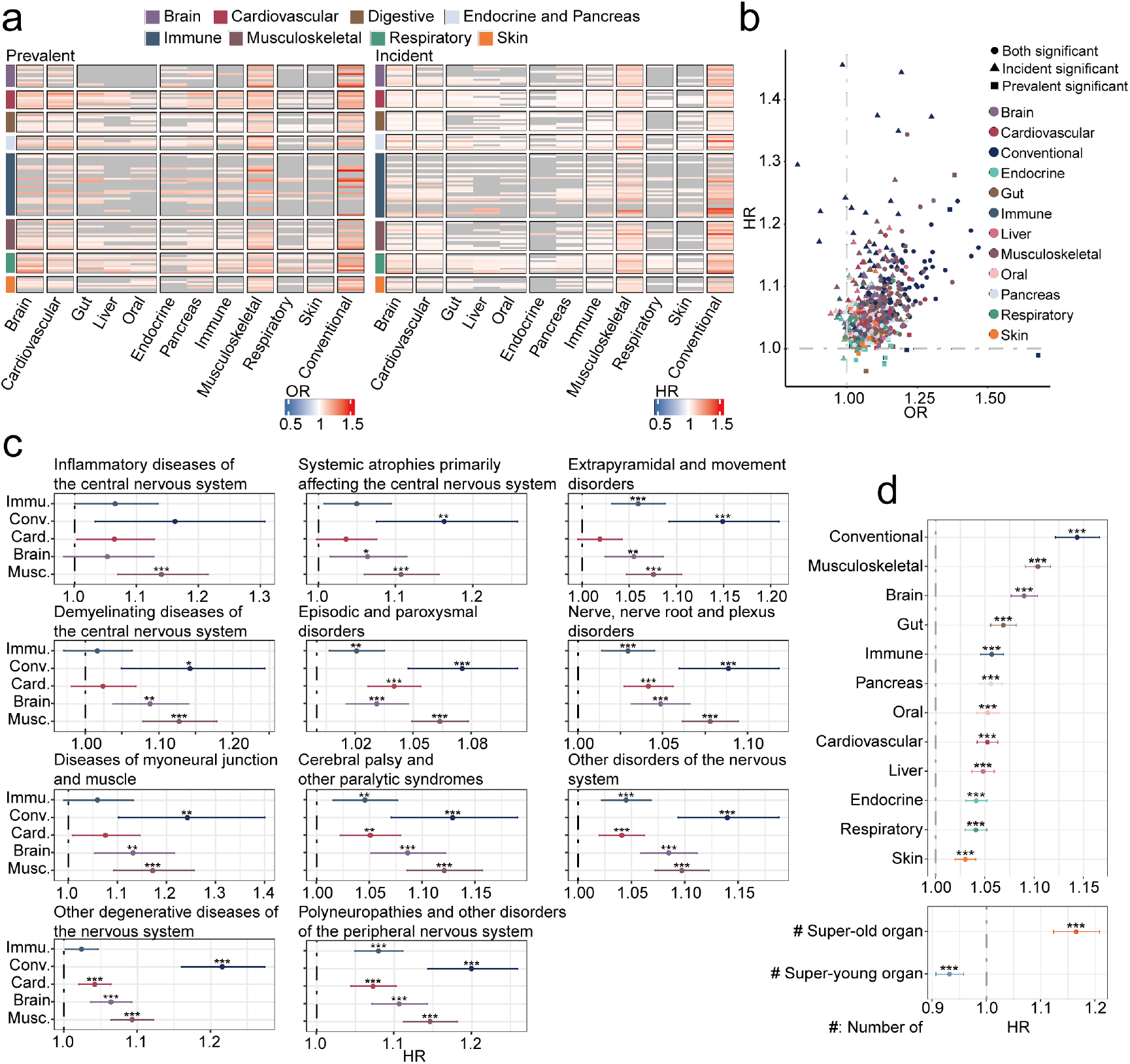
Comprehensive disease-aging relation landscape. **a**. The vast majority of diseases were associated with accelerated aging across multiple organs, observed in both prevalent (left panel) and incident (right panel) cases. **b**. The direction of disease-aging relation was consistent between prevalent (x-axis) and incident (y-axis) cases. **c**. Besides Brain, accelerated aging in the Conventional, Musculoskeletal, Immune, and Cardiovascular systems was also linked to an increased future incidence of nervous system diseases. **d**. Cox proportional hazards regression showed that acceleration in any aging clock was associated with elevated all-cause mortality, with Convention ranks the highest followed by Musculoskeletal (Upper panel). An increase in the number of accelerated-aging organs correlated with higher mortality, whereas younger-organ phenotypes correlated with reduced risk (Lower panel).

Given the pronounced changes in Brain-cAgeDiff observed across disease states (Fig. 4a, b), we specifically examined diseases within the ‘Diseases of the nervous system’ chapter. The Cox proportional hazards regression results showed that each additional year of aging inferred from the Musculoskeletal, Brain, or Conventional clocks significantly elevated the risk of nervous system diseases (Fig. 4c).

We further investigated the association between organ aging rates and all-cause mortality. Among the 12 organ-proxy clocks, the Conventional (HR = 1.14, 95% CI: 1.12-1.17), Musculoskeletal (HR = 1.10, 95% CI: 1.09-1.11), and Brain (HR = 1.09, 95% CI: 1.08-1.10) clocks exhibited the highest hazard ratios for mortality (Fig. 4d). To assess the cumulative impact of extreme aging phenotypes, individuals with an |Organ-cAgeDiff| > 2 SD were categorized as having ‘super-old’ or ‘super-young’ organs. We observed a dose-response relationship between the number of aged organs and survival: each additional ‘super-old’ organ was associated with a 16% increase in mortality risk (HR = 1.16, 95% CI: 1.12-1.21), whereas each additional ‘super-young’ organ conferred a 7% reduction in risk (HR = 0.93, 95% CI: 0.91-0.96) (Fig. 4d).

## Discussion

This study establishes refined plasma proteome–based organ aging clocks by incorporating organ-specific proteome information, advancing beyond previous approaches that largely relied on bulk transcriptomic markers and focused on a narrow spectrum of diseases. By integrating multi-level molecular evidence and data across platforms, our clocks demonstrated enhanced biological interpretability, clinical relevance, and the ability to reveal systemic patterns of aging. In particular, our work extends the scope of prior proteomic aging studies by providing a comprehensive disease–aging landscape.

Several limitations of this study warrant consideration. First, while we integrated multi-omics datasets to identify organ-enriched markers, the representativeness of these proteins in plasma as proxies for their respective organs remains to be further refined. Second, although our study incorporated data from two major proteomic platforms (Olink and SomaScan), the resulting aging clocks are currently not directly interchangeable. The technical biases inherent in different affinity-based assays and the challenge of missing features remain significant hurdles for the clinical application of these models. Overcoming this limitation will require harmonization strategies or multi-platform training to ensure generalizability and translational utility.

## Methods

### Cohort

The UK Biobank is a prospective population-based cohort that collected omics and phenotypic data from approximately 500,000 participants aged 40-69 at recruitment between 2006 and 2010 across the United Kingdom. Detailed descriptions of all available phenotypes can be accessed via the link. All participants provided informed consent.

The UK Biobank Pharma Proteomics Project (UKB-PPP) consortium performed proteomic profiling on plasma samples from 54,219 participants obtained at baseline, using the Olink Explore platform, which quantified 3,072 proteins.

Validation datasets based on the SomaScan platform were derived from demo data provided in the Organage package^16^, and a cohort of 18 healthy controls and 18 patients with Alzheimer’s disease (AD)^23^, as well as a cohort comprising 32 healthy controls and 73 patients with COVID-19^22^. The COVID-19 cohort was assayed using the SomaScan v4 platform, while the remaining cohorts were profiled with version v4.1.

IPF cohort were obtained from a Phase II clinical trial for a novel therapeutic agent^41^. This cohort enrolled 71 patients diagnosed with IPF, who were assigned to four groups including a placebo group and three different dosage experimental groups. Proteomic profiling was conducted using the Olink platform for 43 volunteers from this cohort.

### Data quality control, curation, and preprocessing

To account for measurement disagreements stemming from technical differences between platforms, z-score normalization and quality control were applied separately to the Olink UKB and SomaScan datasets before any cross-platform comparisons or ensemble modeling.

For UKB cohort, data with an excessive proportion of missing values were first excluded. Specifically, any protein that was undetectable in more than 20% of the participants was classified as a high-missingness protein and removed from the dataset. Similarly, any individual with more than 20% of proteins undetected in their plasma sample was considered a low-quality sample and excluded. After this filtering process, data from 44,190 volunteers and 2,911 proteins were retained for subsequent analysis.

According to the Olink platform specifications, the provided measurements represent Normalized Protein Expression (NPX) values, which reflect relative protein abundance on a log_2_ scale calibrated against internal controls. Since missing values indicate protein concentrations below the detection limit, imputation with zero is not appropriate in the log-transformed space. Therefore, NPX values were first exponentiated (base 2) to revert to a linear scale, after which missing values were imputed with zero.

For data generated using the SomaScan platform, version v4.1 data were converted to the v4 standard using a conversion factor list provided by the manufacturer. When multiple measurements corresponded to the same protein, the average expression value was taken. As per the platform’s documentation, SomaScan data undergo Adaptive Normalization by Maximum Likelihood (ANML). These values were log_10_-transformed prior to subsequent analysis.

### Organ enriched marker identification

We initially selected proteins classified as highly expressed in specific organs according to the HPA. Leveraging advances in single-cell technology, we then obtained cell-specific markers that were uniquely expressed in particular tissue from the CM2 database. Concurrently, we incorporated the GTEx dataset as a complementary resource at the bulk RNA level. Genes showing at least four-fold higher expression in one organ compared to any other organ were designated as organ-specific markers, by following a previously used definition^24^. For each candidate marker derived from any single dataset, it was definitively classified as a marker only if it was not associated with any other organ in both the HPA and CM2 databases.

### Identification of age-associated proteins

For the initial identification of age-associated proteins, to ensure a balanced representation of age and sex across the study population, we selected a stratified subset of 6,200 volunteers (100 males and 100 females for each year from 40 to 70). The sample size of 100 per subgroup was determined based on the smallest demographic subgroup in the overall cohort (females aged 70, n=141), resulting in a sampling proportion of approximately 70%. This approach also had the practical benefit of reducing computational cost and processing time.

For the identification of linearly age-associated proteins, the average protein expression level was first calculated for individuals at each age point. The PCC was then computed between the mean protein expression and age, with significance adjusted using the Benjamini-Hochberg (BH) correction. Proteins with a p-value < 0.05 and an absolute PCC greater than 0.4 were defined as significantly linearly age-associated. To identify non-linearly age-associated proteins, the Fuzzy C-Means Clustering (FCM) algorithm was employed, implemented via the Mfuzz package (v2.52.0). To ensure comparability across time points, both the UK Biobank and SomaScan cohorts were restricted to individuals aged 40–70 years. For robustness, age was binned into 5-year intervals. Mean protein expression was calculated within each bin. The number of clusters was manually set to 4. Only proteins showing a strong association with their assigned cluster (membership > 0.6) were retained for further analysis.

### Construction of PAOPAC

For the UK Biobank cohort, the total of 44,190 volunteers were randomly divided into training and testing sets at an 8:2 ratio. Prior to model construction, all data were standardized. A conventional aging clock was built using all plasma proteins as initial input. In our work, we adopted a probabilistic inference strategy based on publicly available tissue-expression atlases: if a protein is highly and preferentially expressed in a particular organ, we assume that, when detected in plasma, it is most likely derived from that tissue. Organ-proxy aging clocks were trained using only the organ enriched marker proteins as input. Models for which the final feature set contained fewer than five proteins were excluded from further analysis. The LASSO, Ridge, and Elastic Net (EN) models were implemented using the glmnet (v4.1-8) and caret (v6.0-94) packages. A 5-fold cross-validation was applied for each training round. For EN and related methods, 100 penalty coefficients (nlambda) were tested. The mean absolute error (MAE) was used as the loss function, and the model with the smallest MAE was selected. Partial Least Squares Regression (PLSR) and k-Nearest Neighbors (KNN) models were implemented using the scikit-learn (v1.5.1) package, with hyperparameter optimization performed via the hyperopt (v0.2.5) package. For PLSR, the number of components was selected from 1 to 20 or the number of overlapping proteins between the organ markers and the platform-detected proteome, whichever was smaller. For KNN, the number of neighbors (k) was optimized between 5 and 50. Five-fold cross-validation was used for both PLSR and KNN, again with MAE as the evaluation metric. LightGBM (LGBM) and XGBoost models were implemented using the lightgbm (v4.5.0) and xgboost (v2.1.1) packages, respectively. Hyperparameter tuning was conducted over the following ranges: number of trees [100, 200, 300, 400], maximum tree depth [3–6], maximum number of leaves [3–63], learning rate (log-uniform between 0.01 and 0.2), and feature sampling ratio [0.6–1.0]. Hyperparameter optimization and feature selection were performed using the shaphypetune (v0.2.7) package, with a maximum of 200 iterations, 20 model trainings per iteration, and a shadow variable ratio of 100%. SHAP values were used to assess feature importance. In preliminary model comparisons and training conducted on the SomaScan dataset, only sex was included as a covariate. For the final aging clocks built on the UKB cohort, the following covariates were included to adjust for potential confounding effects: sex, ethnicity, blood sample collection time, fasting time before sample collection, body mass index (BMI), and Townsend Deprivation Index (TDI). Sex (Field 31) was coded as female = 1, male = 0. Ethnicity (Field 21000) was coded as White = 0, other = 1. Blood sample collection season (Field 3166) was binarized as winter/spring (December–May) = 0 and summer/autumn (June–November) = 1. Fasting time (Field 74), BMI (Field 23104), and TDI (Field 22189) were used as continuous variables after standardization.

### Calculation and adjustment of biological age

The age predicted by the aging clocks is referred to as the biological age (BA). The difference between this BA and the chronological age (CA) is defined as the Age Difference (AgeDiff), which reflects an individual’s rate of aging. A higher AgeDiff indicates accelerated aging, while a lower value suggests decelerated aging. To correct for bias introduced during model training, locally estimated scatterplot smoothing (LOESS) regression was applied to model the relationship between AgeDiff and CA. The fitted values from this regression represent age-related prediction bias (AgeDiff hat). The corrected biological age (cBA) was obtained by subtracting AgeDiff hat from the BA. The corrected Age Diff (cAgeDiff) was then derived by subtracting the CA from cBA. Individuals with an absolute cAgeDiff greater than two standard deviations were classified as either ‘super-younger’ or ‘super-older’.

### Acquisition of event outcomes

Event outcomes and their first occurrence dates in the UK Biobank cohort were extracted using Fields 41270 and 41280, respectively. Disease codes were mapped to specific clinical conditions based on the ICD-10 classification system. Diseases first reported prior to enrollment in the cohort were classified as prevalent cases, while those newly identified during follow-up visits were classified as incident cases. Individuals with no recorded disease events throughout the study period were considered completely healthy controls. Date of death was obtained from Field 40007. The last follow-up time or loss-to-follow-up date was retrieved from Field 20143.

### Pathway enrichment analysis

Pathway enrichment analysis was performed using the clusterProfiler package (v4.0.5). Gene Ontology Biological Process (GO BP) terms were used as the reference gene set. A significance cutoff of 0.05 was applied, and the Benjamini–Hochberg (BH) method was used for multiple testing correction. In visualizations of results, the term ‘p-value’ refers to the original, unadjusted significance value; all other reported values are adjusted. Reduction and visualization of overlapping GO terms were carried out with the rrvgo package (v1.4.4), using the GO BP annotation set. Similarity between child terms was computed based on the ‘Rel’ semantic similarity measure, with a similarity threshold set to 0.7 for term clustering and simplification.

## Data availability

To request access of UK Biobank, all bona fide researchers who wish to conduct health-related research can apply for access to UK Biobank through UK Biobank’s access management system. Organage demo data is available at Github (https://github.com/hamiltonoh/organage/tree/main/tests). AD data is available at Synapse (https://www.synapse.org/Synapse:syn30549757/files/). COVID data is available at Mendeley Data (https://data.mendeley.com/datasets/2mc6rrc5j3/1). IPF data is available at OMIX by OMIX008341.

## Code availability

PAOPAC models are publicly available at. https://github.com/JackieHanLab/PAOPAC.

---

**Supplementary Figure 1.**
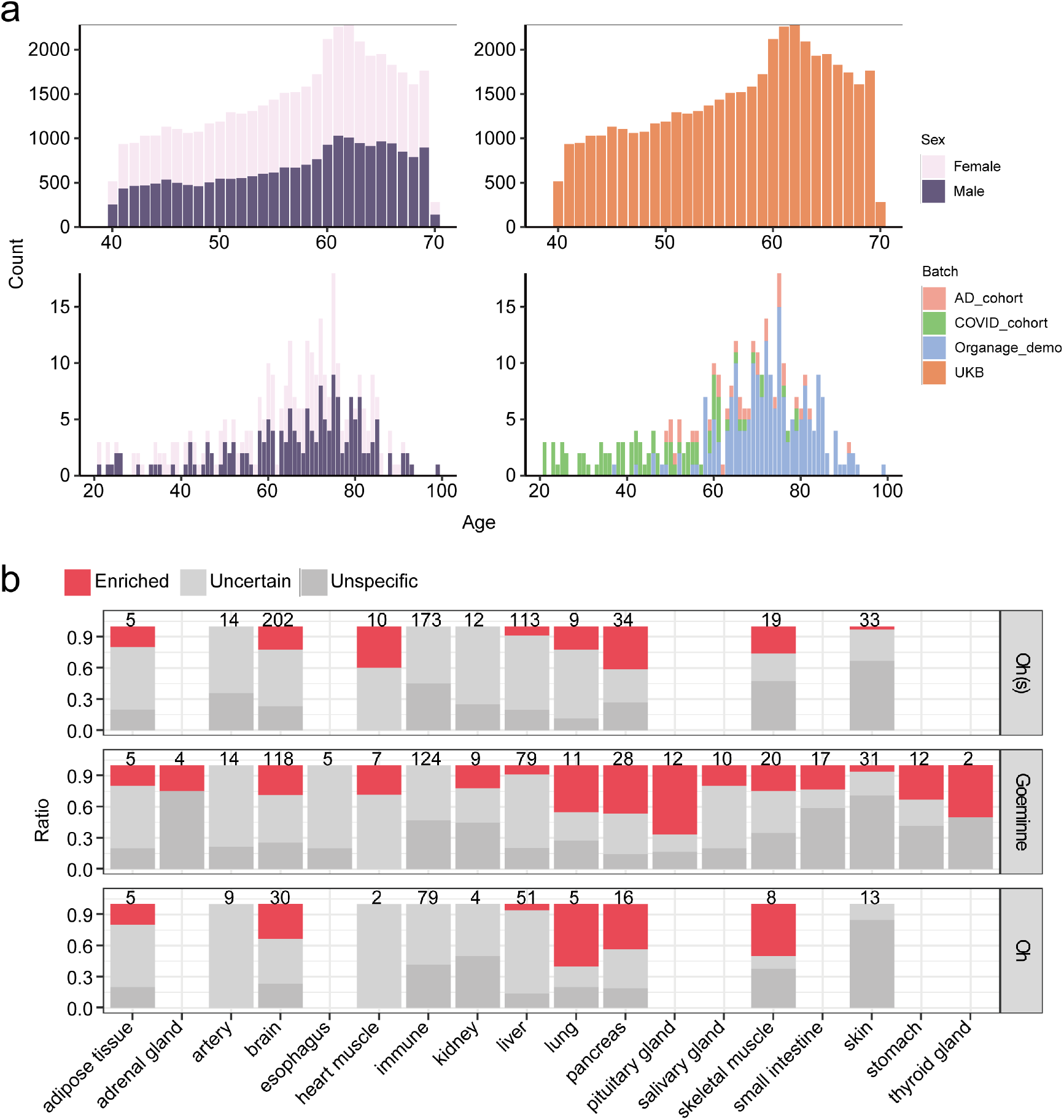
Description of datasets and validation of organ-enriched markers. **a**. Distribution of age, sex, and batch across the datasets. **b**. At least 20% of organ-enriched markers in previous studies are unspecific against the Human Protein Atlas (HPA). ‘Enriched’ indicates that a protein is highly expressed in only one organ; ‘Unspecific’ refers to proteins with high expression in multiple organs; the remaining are classified as ‘Uncertain’. Oh (s) denotes the published model trained on SomaScan data, the other two models were trained on Olink data. HPA, Human Protein Atlas.

**Supplementary Figure 2.**
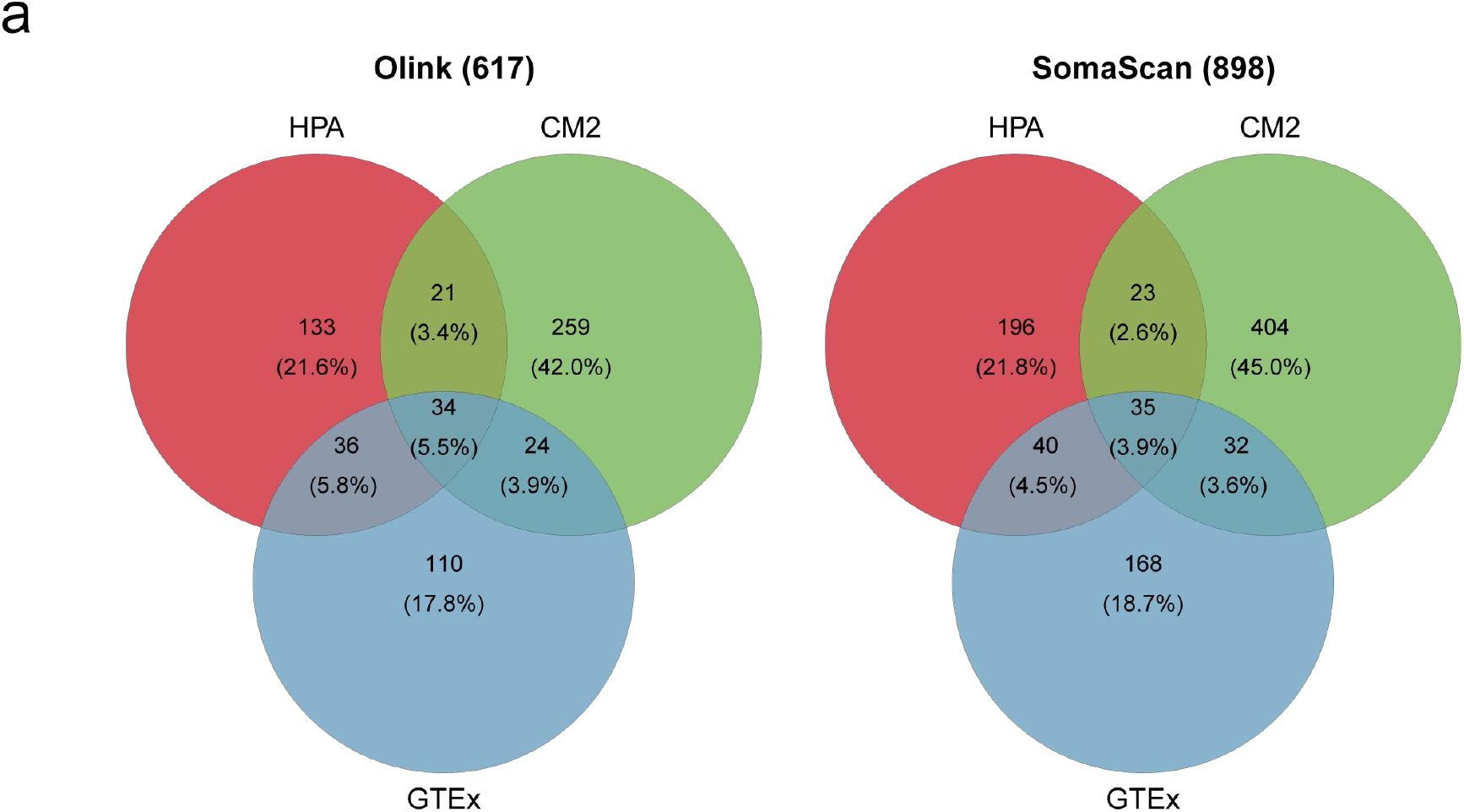
Organ-enriched proteins’ overlap across proteomic platforms. **a**. Venn diagrams illustrating the overlap of organ-enriched proteins defined by three datasets (HPA, CM2, and GTEx) in the Olink and SomaScan proteomic platforms, with the number and percentage of proteins in each segment indicated.

**Supplementary Figure 3.**
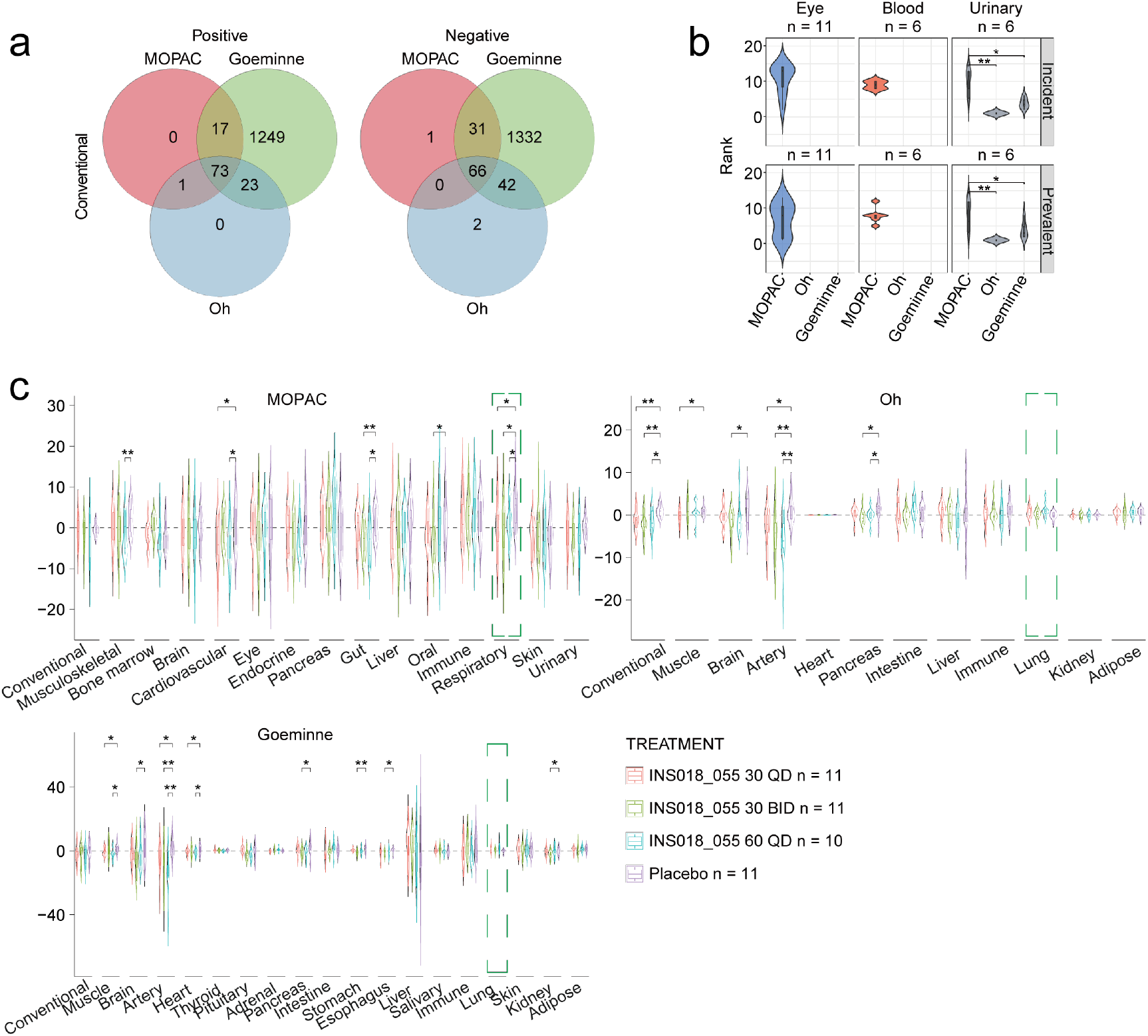
Comparison between PAOPAC and previous work. **a**. Venn diagrams illustrating the overlap of features utilized in the Conventional aging clock across three distinct studies. Owing to the application of more stringent feature selection criteria, PAOPAC retained the smallest number of features, and its feature set constituted a strict subset of those employed in the other two studies. **b**. While PAOPAC incorporates two novel organ-proxy clocks, their overall performance is moderate. And the Urinary clock underperforms relative to the previously published clocks. **c**. Changes in biological age following treatment in IPF patients, as assessed by different aging clocks. Only PAOPAC in this study tells the delayed Respiratory aging in treatment group (t-test). The Respiratory or Lung clock is labelled by green box. IPF, idiopathic pulmonary fibrosis.

